# Intrinsic variation in the vertically transmitted insect-specific core virome of *Aedes aegypti*

**DOI:** 10.1101/2021.08.30.458191

**Authors:** H. Coatsworth, J Bozic, J. Carrillo, E.A. Buckner, A.R. Rivers, R.R. Dinglasan, D.K. Mathias

## Abstract

Since 2009, local outbreaks of dengue (serotypes 1-3) mediated by *Aedes aegypti* mosquitoes have occurred in the United States, particularly in Florida (FL). In 2016 and 2017, dengue virus serotype 4 was found alongside several insect-specific viruses (ISVs) in pools of *A. aegypti* from sites in Manatee County, FL, in the absence of an index case. Although ISVs have been characterized in *A. aegypti* globally, the constitution of a core virome in natural populations remains unclear. Using mosquitoes sampled from the same area in 2018, we compared baseline ovary viromes of field (G_0_) and lab (Orlando) *A. aegypti* via metagenomic RNA sequencing. Across all samples, virome composition varied by sample type (field- or colony-derived). Four ISVs comprised >97% of virus sequences: a novel partiti-like virus (Partitiviridae), a previously described toti-like virus (Totiviridae), unclassified Riboviria, and four previously described orthomyxo-like viruses (Orthormyxoviridae). Whole or partial genomes for the toti-like virus, partiti-like virus, and one orthomyxo-like virus were assembled and analyzed phylogenetically. Multigenerational maintenance of these ISVs was confirmed orthogonally by RT-PCR in G_0_ and G_7_ mosquitoes, indicating vertical transmission as the mechanism for ISV sustentation. This study provides fundamental information regarding ISV ecology, persistence, and variation in *A. aegypti* in nature.

## Introduction

Mosquito-borne viral diseases such as dengue, chikungunya, and Zika have spread to new areas, putting over half the world’s population at risk of infection (Weaver 2014, Kraemer et al. 2015). This increase is largely correlated with the global expansion of *Aedes aegypti* and *Aedes albopictus* mosquitoes, the primary vectors for these arboviruses (Messina et al. 2019, Brady and Hay 2020). In Florida, USA, both vector species are present, and *A. aegypti* populations thrive in most of the state’s large population centers due to their urban niche (Britch et al. 2008, Reiskind and Lounibos 2013, Wilke et al. 2019) and widespread insecticide resistance (Estep et al. 2018, Mundis et al. 2020).

Autochthonous dengue virus (DENV, serotypes 1-3) infections have occurred in 11 of the last 12 years in urban centers in South Florida (CDC 2010, Rey 2014, https://www.cdc.gov/dengue/statistics-maps/index.html), while in 2014 and 2016 outbreaks of chikungunya (Kendrick et al. 2014) and Zika (Likos et al. 2016, Grubaugh et al. 2017), respectively, occurred exclusively in counties with recent permanent population records of *A. aegypti* (Hahn et al. 2017). Since the re-emergence of DENV in the Florida Keys in 2009, local transmission of at least one serotype has occurred in the state each year except in 2017. Moreover, despite the absence of a local human index case, dengue virus serotype 4 (DENV-4) was detected and fully sequenced alongside numerous insect-specific viruses (ISVs) in the abdomens of *A. aegypti* adult G_0_ females from Manatee County, Florida, in 2016 and 2017 (Boyles et al. 2020).

The ability of a mosquito to transmit DENV (i.e., its vector competence) varies based on external environmental variables, the genetic backgrounds of the vector and virus, and the composition of the mosquito’s microbiome (Tabachnick 2013). The latter is complex and includes ISVs, which can only replicate in the cells of insect hosts (Sang et al. 2003, Nasar et al. 2012, Junglen et al., 2017). ISVs are pervasive in both wild-caught and laboratory-reared mosquitoes, and co-infection of ISVs from the same viral family as DENV are known to influence the vector competence of *A. aegypti* (Bolling et al. 2012, Zhang et al. 2017, Atoni et al. 2019, Baidaliuk et al. 2019, Öhlund et al. 2019). However, most recently discovered ISVs belong to other virus families, and given their pervasiveness, a better understanding of their ecology and impact on mosquito biology is needed. The term “core virome” was recently coined to describe the set of ISVs common to the majority of individuals in a mosquito population (Shi et al. 2019). The term has since been divided into two categories: vertical (passed from mother to offspring) and environmental (acquired from the environment) (Shi et al. 2020). To date, research on *Aedes* mosquitoes suggests that at the population level core viromes are maintained across developmental stages (Shi et al. 2020) and over short time scales (∼ 1 year) (Boyles et al. 2020, Shi et al. 2020). However, it is unknown how well vertical core viromes are sustained over longer periods of time, or the degree to which environmental core viromes are transient. Moreover, most colonies commonly used in arbovirus research have yet to be examined, and these could be useful models to address questions about the impact of core viromes on mosquito biology.

To begin filling these gaps in knowledge, we used tissue-specific RNA metagenomics and reverse-transcription PCR to follow up on the previous virome profiles characterized for Floridian *A. aegypti* (Boyles et al. 2020). Our objectives were to i) define the vertically transmitted core virome in ovary pools from G_0_ field- and lab-derived *A. aegypti*, ii) examine the persistence of specific core ISVs over time in ovaries of G_7_ descendants of field-derived females, iii) assess variability among the sexes for core ISVs maintained in a field-derived colony, and iv) use phylogenetics to investigate evolutionary relationships between core virome members and previously described ISVs. We hypothesized that elements of the vertical core virome would be present in both field- and lab-derived “Orlando” (ORL) strain *A. aegypti* due to both originating from Florida, but that field-derived samples would have higher ISV diversity due to greater environmental variability. We further predicted that male ISV infection status would mimic that of their female counterparts because of efficient vertical transmission and that *A. aegypti* ISVs would cluster phylogenetically with ISVs from other mosquito taxa.

## Results

### Partiti- and toti-like insect-specific viruses dominate the *Aedes aegypti* virome

To identify core virome ISV members as well as compare field versus lab *A. aegypti* virome diversity, viral reads from each ovary pool were assigned to their lowest common ancestor (LCA) and summed based on read count. Partitiviridae reads comprised between 50-76% of all viral reads in the field-derived Manatee County (Palmetto, P) samples, while unclassified Riboviria reads made up >57% of the viral reads in the ORL sample and between 5-11% of the P samples (Figure 1). Viral reads aligning best with Atrato partiti-like virus 3 were the most prevalent across all the P samples, making up 70, 72, 51, 67 and 76 % of the viral reads (P1 – 5 ovary pools respectively), but were completely absent from the ORL sample. Reads aligning to *A. aegypti* toti-like virus accounted for 19, 17, 33, 10, and 11% of the reads for the P samples, and represented 42% of the ORL sequences. Together, 75-90% of all viral reads from field-derived samples aligned to these two groups (Partitiviridae and Totiviridae). In addition, 42% of the ORL reads and ∼5% of reads in each of the P samples aligned with an unclassified Riboviria virus, followed by 13% matching to a dsRNA virus environmental sample in ORL and ∼1.5% in all P samples. Sequences matching to four viruses in Orthomyxoviridae (Guadeloupe mosquito quaranja-like virus 1, Whidbey virus, unclassified Orthomyxoviridae, and *Aedes alboannulatus* orthomyxo-like virus) were restricted to field-derived samples and collectively accounted for 1.7, 2.4, 2.0, 11.4, and 7.2% of the reads from P1 – P5, respectively. The remaining viral sequences in these samples each represented less than one percent of the total viral sequence pool, and in order of prevalence, matched to *A. aegypti* virga-like virus, *A. aegypti* anphevirus, unclassified viruses, Chuvirus Mos8Chu0, Liao ning virus, and an unclassified flavivirus (Figure 1). The field-derived P samples were more diverse than the ORL sample, as the P samples had an average of 12 different viral assignments, while the ORL sample only contained 5.

**Figure 1.**
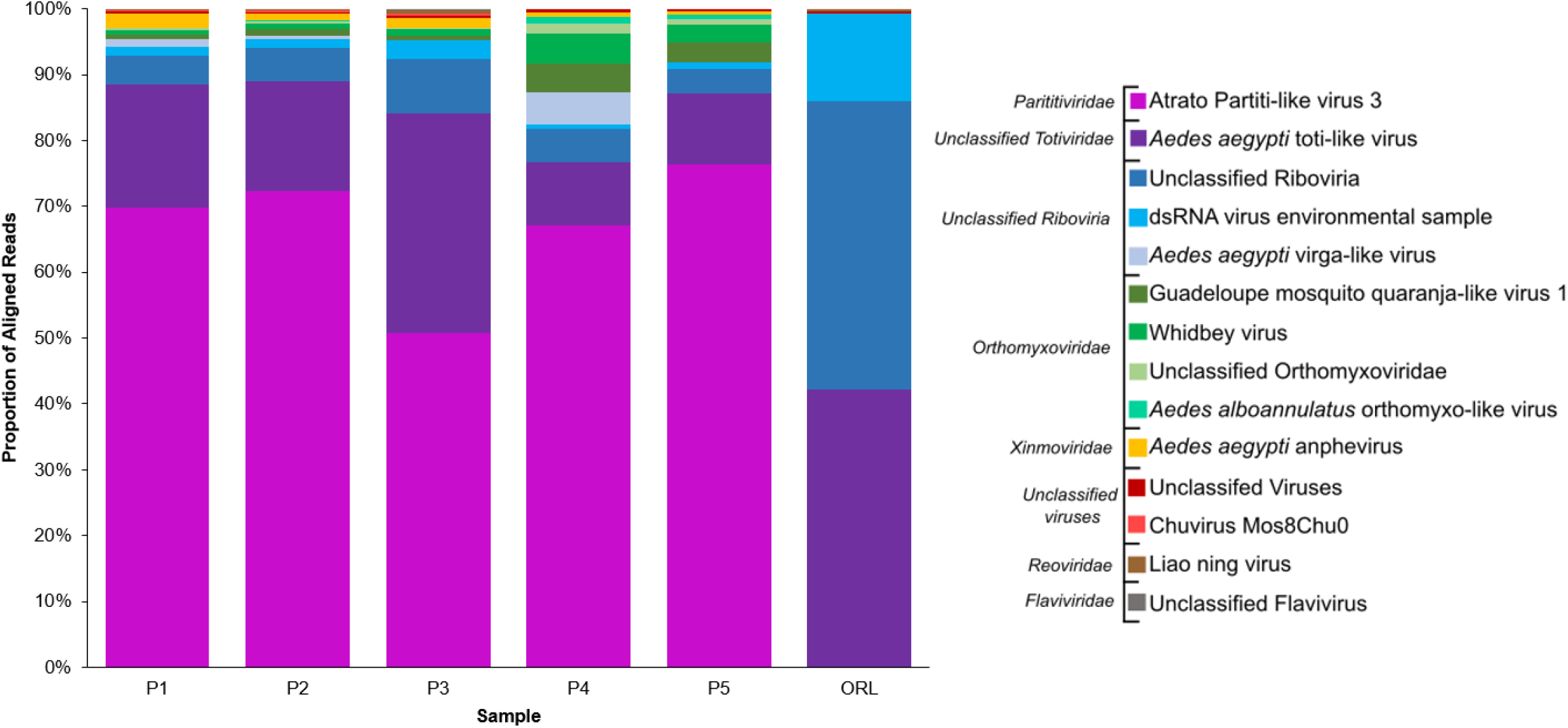
Proportion of aligned viral reads across ovary pools of field-derived Palmetto (P, Manatee Country, FL, USA) and lab-reared Orlando (ORL) *A. aegypti*. The samples are ranked in descending order of viral read prevalence, grouped, and colored by viral classification. Fuchsia: Partitiviridae, purple: unclassified Totiviridae, blue: unclassified Riboviria, green: Orthomyxoviridae, yellow: Xinmoviridae, red: unclassified viruses, brown: Reovirdiae, grey: Flaviviridae.

### Genome assembly of Palmetto toti-, partiti- and orthomyxo-like viruses

Whole or partial genome assembly was completed for our Palmetto toti-, partiti- and orthomyxo-like viruses to compare the overall similarity between the viruses found here and those reported previously. Eight open reading frames (ORFs) were predicted based on the assembled Palmetto toti-like virus contigs. Only 2 of these ORFs (ORF 2 [1,033aa] and ORF 3 [1,021aa]) assembled in a linear format akin to the typical capsid-RdRp *Totiviridae* structure (Figures 2, 3) and created products with known sequence similarity. The 1,033aa long ORF of the Palmetto toti-like virus showed high amino acid similarity to other *A. aegypti* RNA-dependent RNA polymerase (RdRp) genes in toti-like viruses (>98% identity), with high query coverage (∼90%) to 926aa RdRp sequences isolated from individual *Aedes aegypti* mosquitoes from Guadeloupe, a Caribbean island (QEM39131.1 and QEM39133.1) (Shi et al., 2019). The 1,021aa sequence showed high similarity and query coverage to the 1,008aa long Guadeloupe *Aedes aegypti* toti-like capsid sequences (>99% and 98%, respectively) isolated from the samples. Together, these sequences represent the RdRp and capsid genes of the Palmetto toti-like virus identified herein.

**Figure 2.**
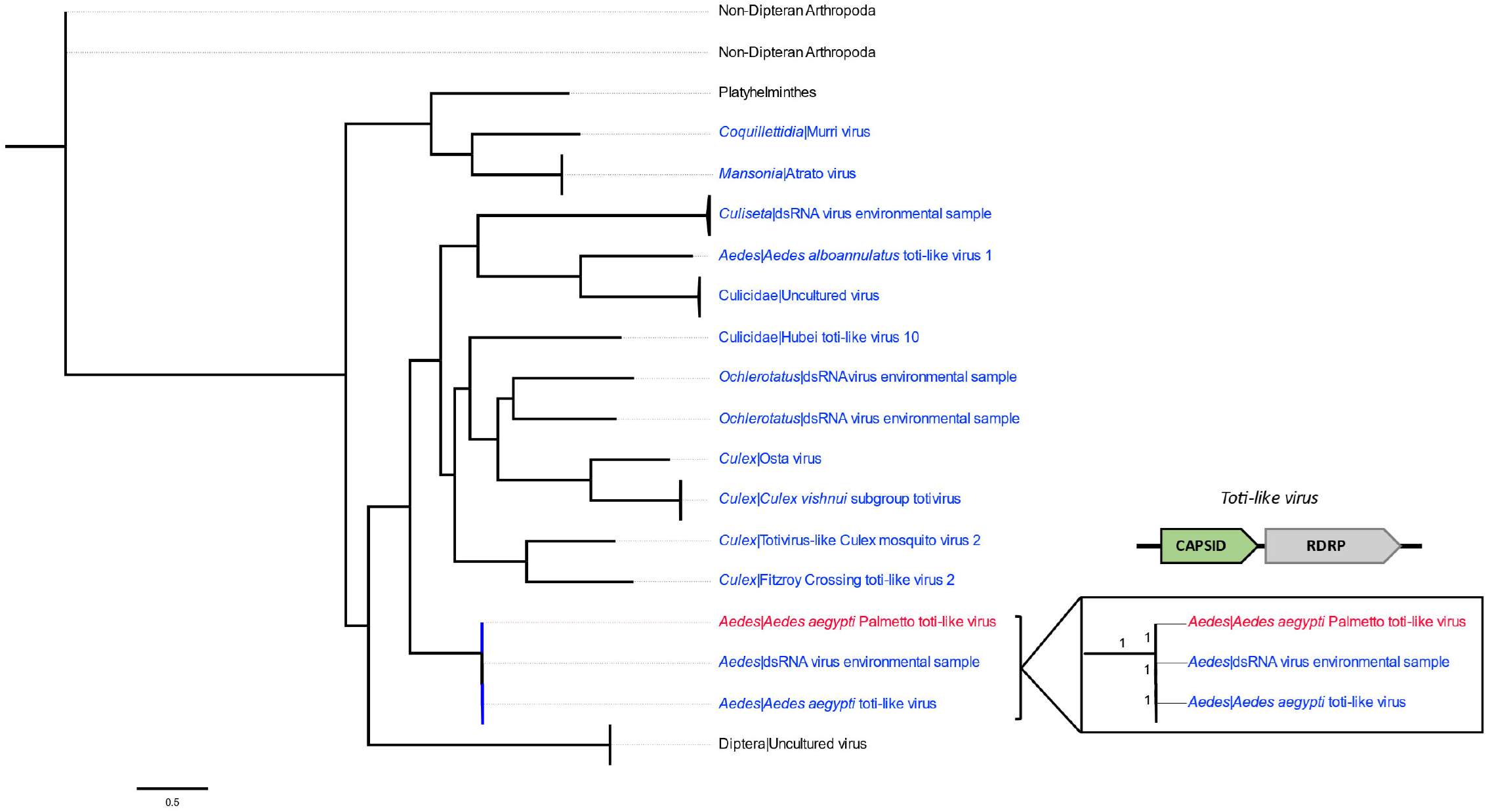
Maximum likelihood tree of toti-like virus capsid sequences (using an LG+G+I+F model) based on amino acid sequences. The tree is representative of 25 sequences spanning 1008 sites and 47 branches. Mosquito-derived virus sequences are colored in blue, while the Palmetto sequence is in red. Virus names are appended to all sequences derived from Diptera. The generalized viral genomic structure is to the right of the tree, with the segment analyzed in color (green polygon represents capsid/nucleoprotein sequence). The numbers on each branch within the inset denote branch support lengths. Scale bar is representative of 0.5 amino acid substitutions per site.

**Figure 3.**
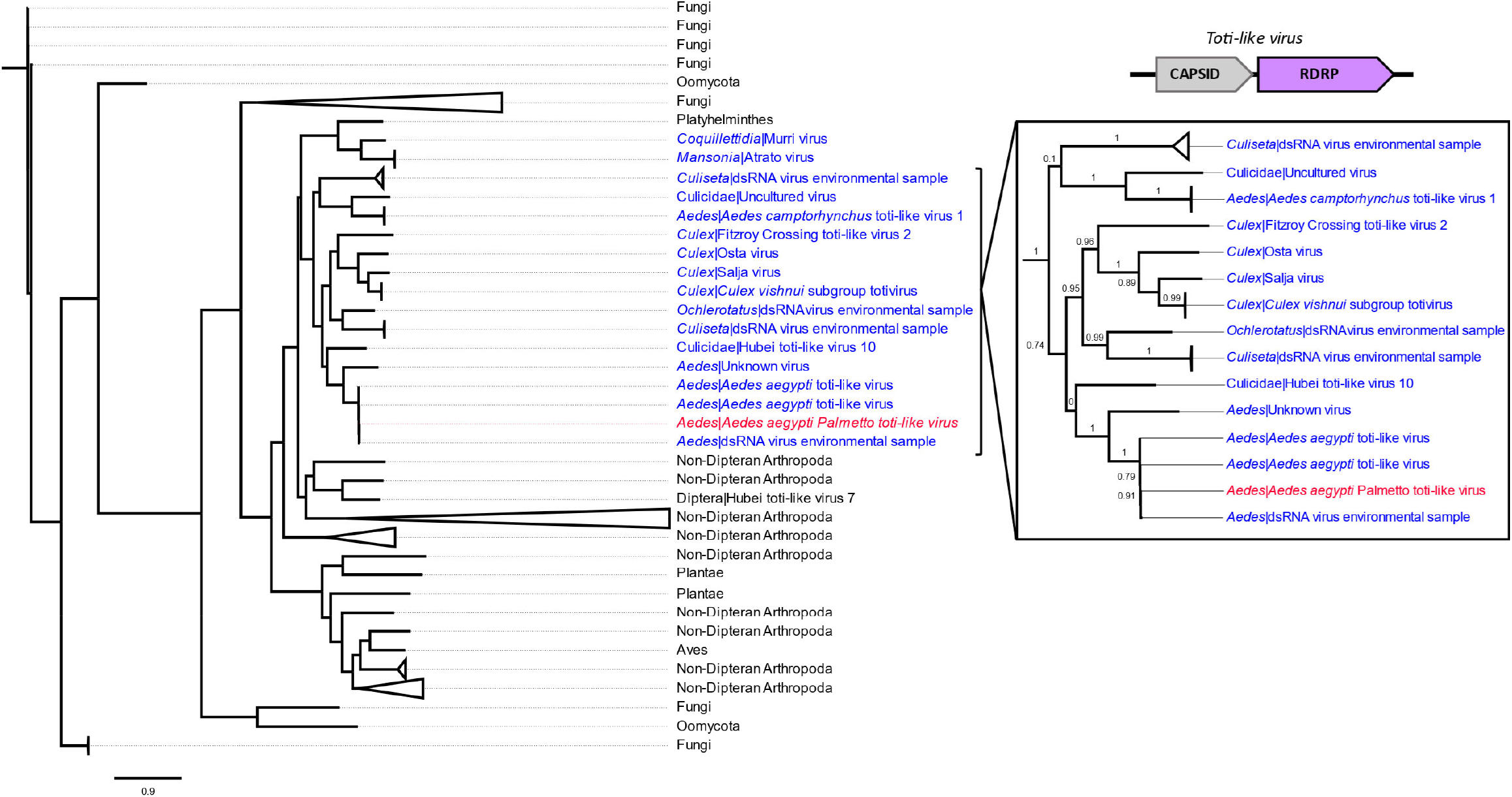
Maximum likelihood tree of toti-like virus RdRp sequences (using an LG+G+I+F model) based on amino acid sequences. The tree is representative of 101 sequences spanning 1032 sites and 199 branches. Mosquito-derived virus sequences are colored in blue, while the Palmetto sequence is in red. Virus names are appended to all sequences derived from Diptera. The generalized viral genomic structure is to the right of the tree, with the segment analyzed in color (purple polygon represents polymerase sequence). The numbers on each branch within the inset denote branch support lengths. Scale bar is representative of 0.9 amino acid substitutions per site.

In contrast, only three ORFs were predicted for the Palmetto partiti-like virus, which likely has a simple two-segmented genomic structure characteristic of the Partitiviridae (Figure 4). Only one of these ORFs (ORF 1 [446aa]) had sequence similarity to known sequences, with 78% identity and 95% query coverage to a 472aa RdRp Atrato partiti-like virus sequence (QHA33899.1) isolated from *Psorophora albipes* mosquitoes in Colombia, and 79% identity and 86% query coverage to another RdRp Atrato partiti-like virus sequence (QHA33901.1, 387aa) isolated from *Anopheles darlingi* mosquitoes in Colombia. Therefore, this sequence was classified as the RdRp for the Palmetto partiti-like virus.

**Figure 4.**
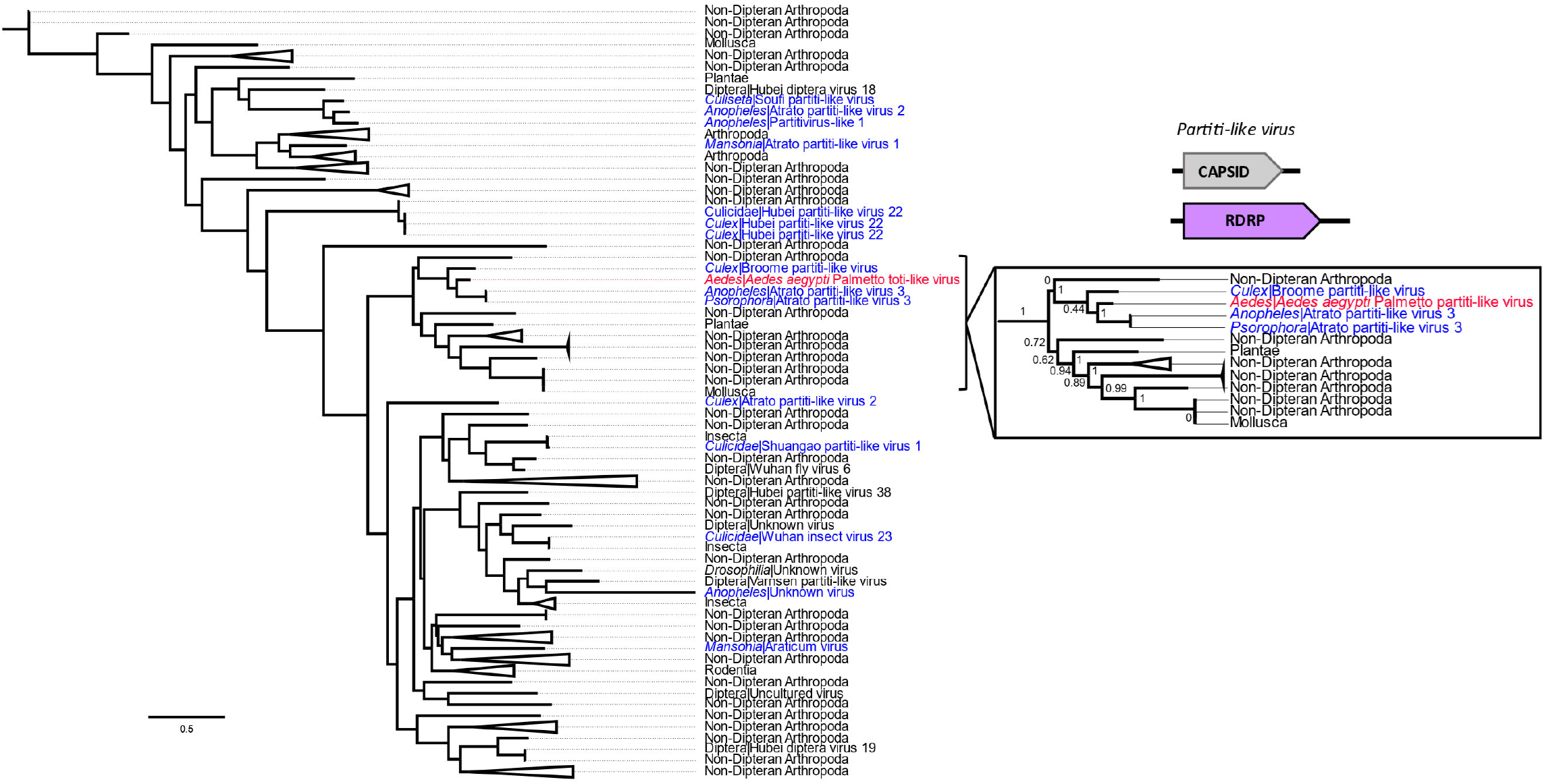
Maximum likelihood tree of partiti-like RdRp virus sequences (using an LG+G+I+F model) based on amino acid sequences. The tree is representative of 101 sequences spanning 447 sites and 199 branches. Mosquito-derived virus sequences are colored in blue, while the Palmetto sequence is in red. Virus names are appended to all sequences derived from Diptera. The generalized viral genomic structure is to the right of the tree, with the segment analyzed in color (purple polygon represents polymerase sequence). The numbers on each branch within the inset denote branch support lengths. Scale bar is representative of 0.5 amino acid substitutions per site.

Twenty ORFs were predicted from the Orthomyxoviridae contigs, and 6 of these ORFs had known sequence similarity based on non-redundant protein blast searches. Four of these ORFs appeared to represent one virus (Palmetto orthomyxo-like virus), while the remaining two ORFs are likely from two other orthomyxoviruses: ORF 4 (229aa) with 33% identity and 65% query coverage to an Atrato Chu-like virus 5 hypothetical protein 2 (QHA33674.1) identified in *Psorophora albipes* from Colombia, and ORF 5 (294aa) with 66% identity and 87% query coverage with the nucleoprotein of Wuhan Mosquito Virus 6 (QRW42410.1) isolated from *Culex tarsalis* mosquitoes from California. The 4 Palmetto orthomyxo-like virus ORFs had highest sequence similarity to Guadeloupe mosquito quaranja-like virus 1 (GMQLV1), isolated from *A. aegypti* collected in San Diego County, California (Batson et al. 2021). Like other viruses in the Orthomyxoviridae, quaranjaviruses have a segmented genome, and GMQLV1 was originally described as likely having 6 or 7 segments (Shi et al.2019). However, recently reported data (Batson et al. 2021) indicate that 8 segments are more likely (Figures 5-8). ORF7 (800aa) had 100% identity and 98% query coverage to GMQLV1 PB2 (QRW42587.1); ORF 10 (797aa) had 99.75% identity and 98% query coverage to GMQLV1 PB1 (QRW42591.1); ORF 11 (716aa) had 99.86% identity and 98% query coverage to GMQLV1 PA (QRW42581.1); and ORF 16 (584aa) had 99.64% identity and 95% query coverage to GMQLV1 NP (QRW42580.1). These 4 ORFs likely represent the nucleoprotein (NP - ORF 16), and the heterotrimeric RdRp complex (PB2 – ORF7, PB1 – ORF10, PA – ORF11) for our Palmetto orthomyxo-like virus, which is likely identical to GMQLV1.

**Figure 5.**
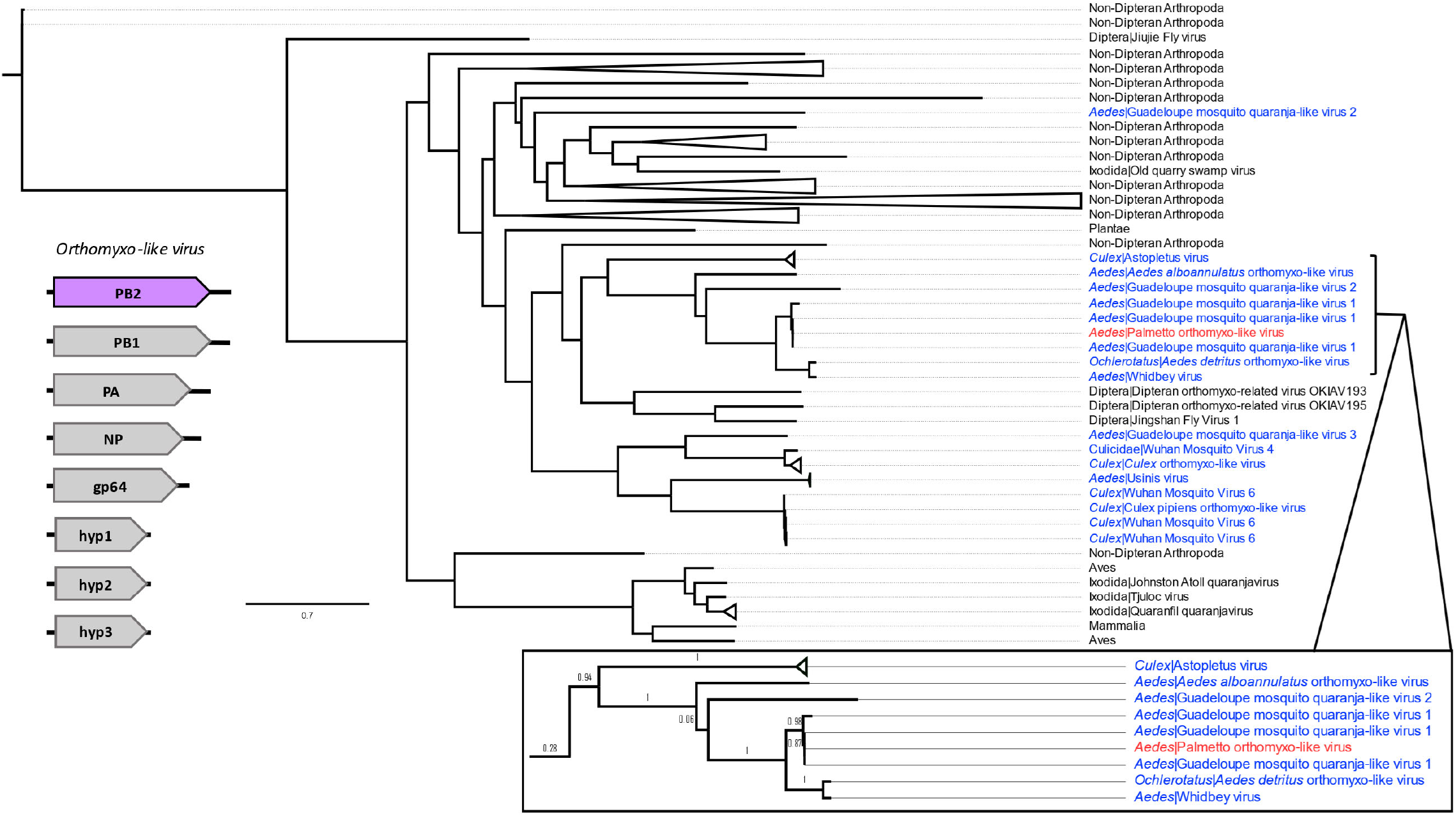
Maximum likelihood tree of orthomyxo-like of the PB2 virus sequences (using an LG+G+F model) based on amino acid sequences. The tree is representative of 78 sequences spanning 1668 sites and 153 branches. Mosquito-derived virus sequences are colored in blue, while the Palmetto sequence is in red. Virus names are appended to all sequences derived from Diptera or Ixodida. The generalized viral genomic structure is to the right of the tree, with the segment analyzed in color (purple polygon represents polymerase sequence). The numbers on each branch within the inset denote branch support lengths. Scale bar is representative of 0.7 amino acid substitutions per site.

**Figure 6.**
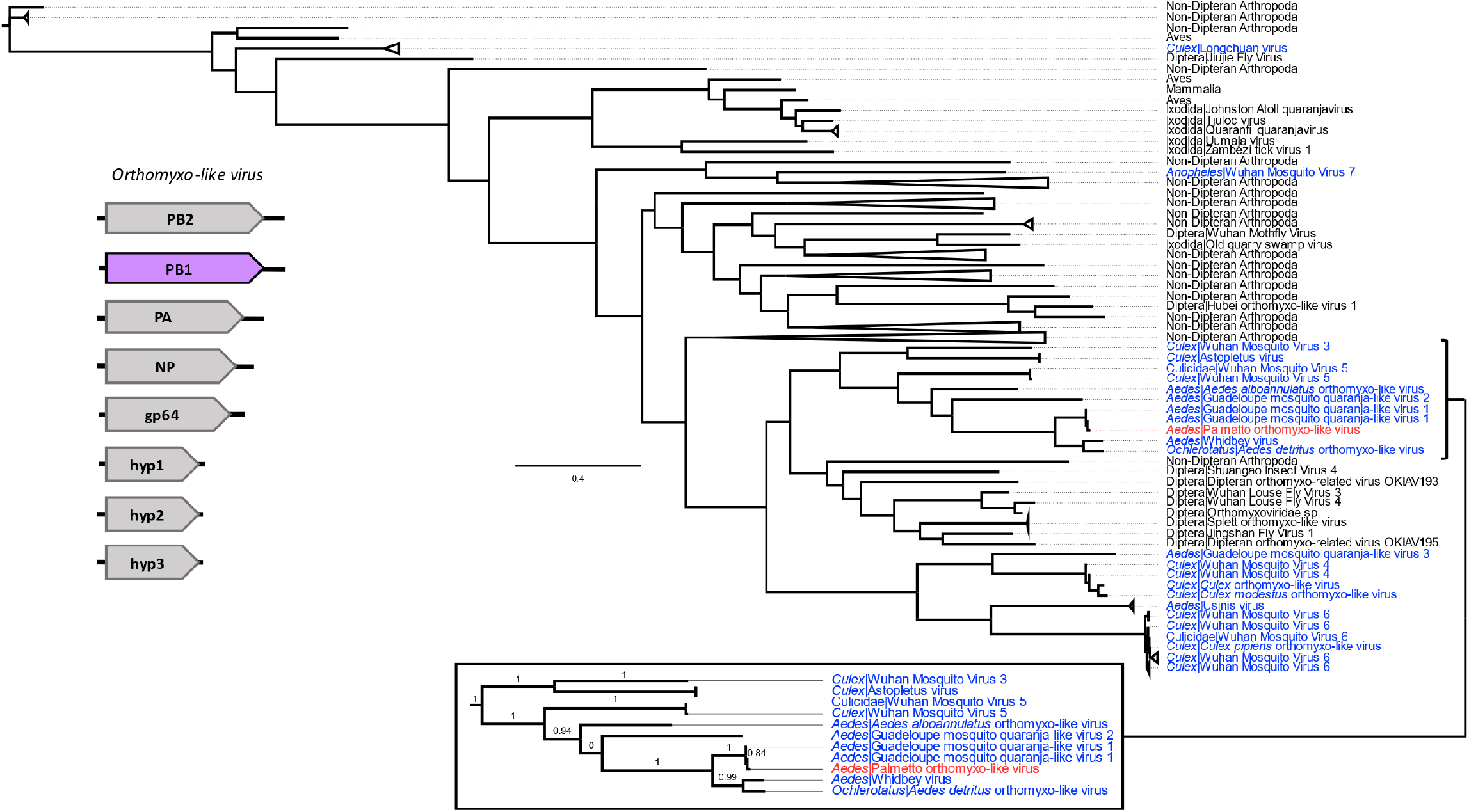
Maximum likelihood tree of orthomyxo-like of the PB1 virus sequences (using an LG+G+I+F model) based on amino acid sequences. The tree is representative of 101 sequences spanning 1041 sites and 199 branches. Mosquito-derived virus sequences are colored in blue, while the Palmetto sequence is in red. Virus names are appended to all sequences derived from Diptera or Ixodida. The generalized viral genomic structure is to the right of the tree, with the segment analyzed in color (purple polygon represents polymerase sequence). The numbers on each branch within the inset denote branch support lengths. Scale bar is representative of 0.4 amino acid substitutions per site.

**Figure 7.**
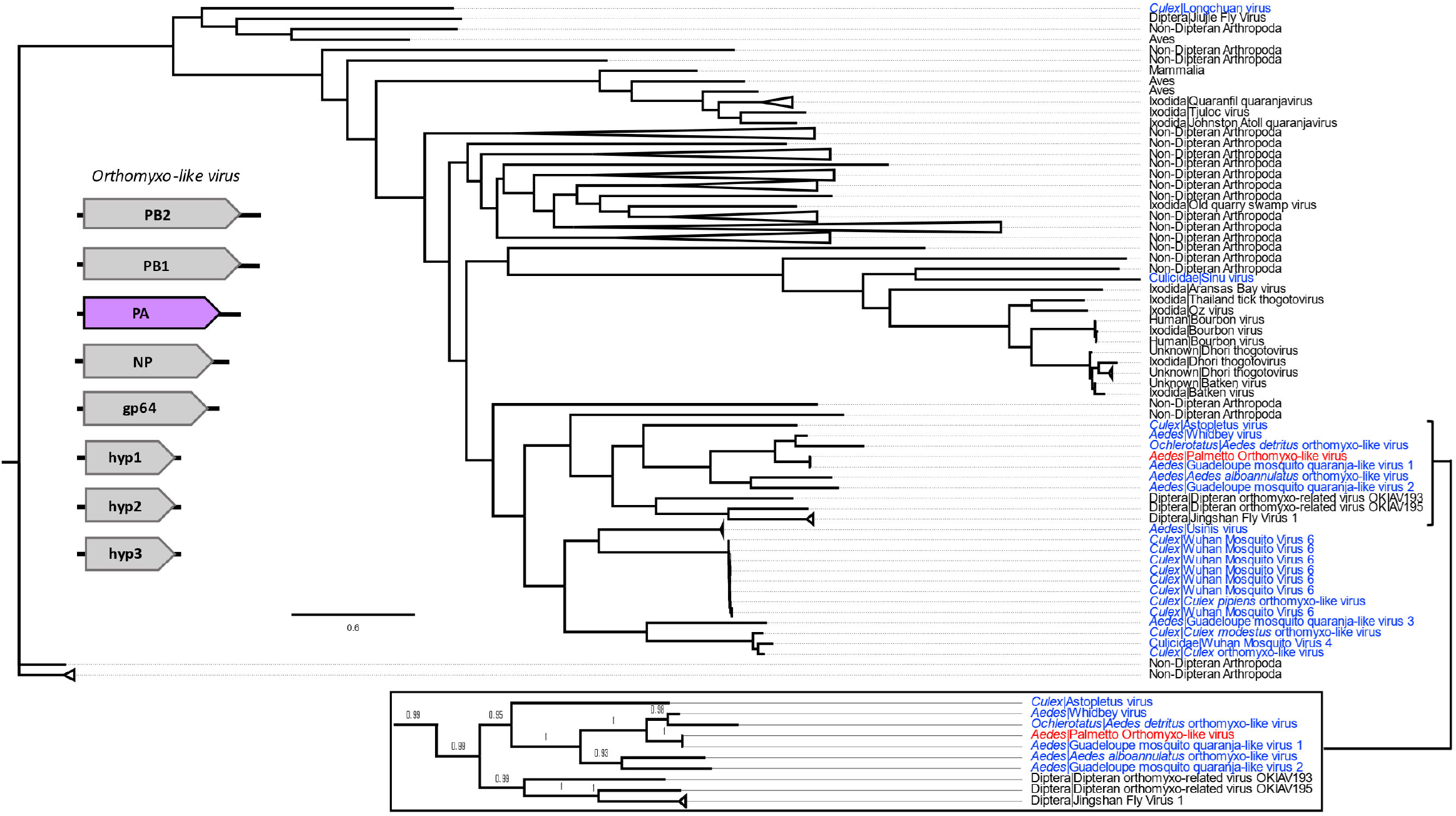
Maximum likelihood tree of orthomyxo-like of the PA virus sequences (using an LG+G+I+F model) based on amino acid sequences. The tree is representative of 101 sequences spanning 1388 sites and 199 branches. Mosquito-derived virus sequences are colored in blue, while the Palmetto sequence is in red. Virus names are appended to all sequences derived from Diptera or Ixodida. The generalized viral genomic structure is to the right of the tree, with the segment analyzed in color (purple polygon represents polymerase sequence). The numbers on each branch within the inset denote branch support lengths. Scale bar is representative of 0.6 amino acid substitutions per site.

**Figure 8.**
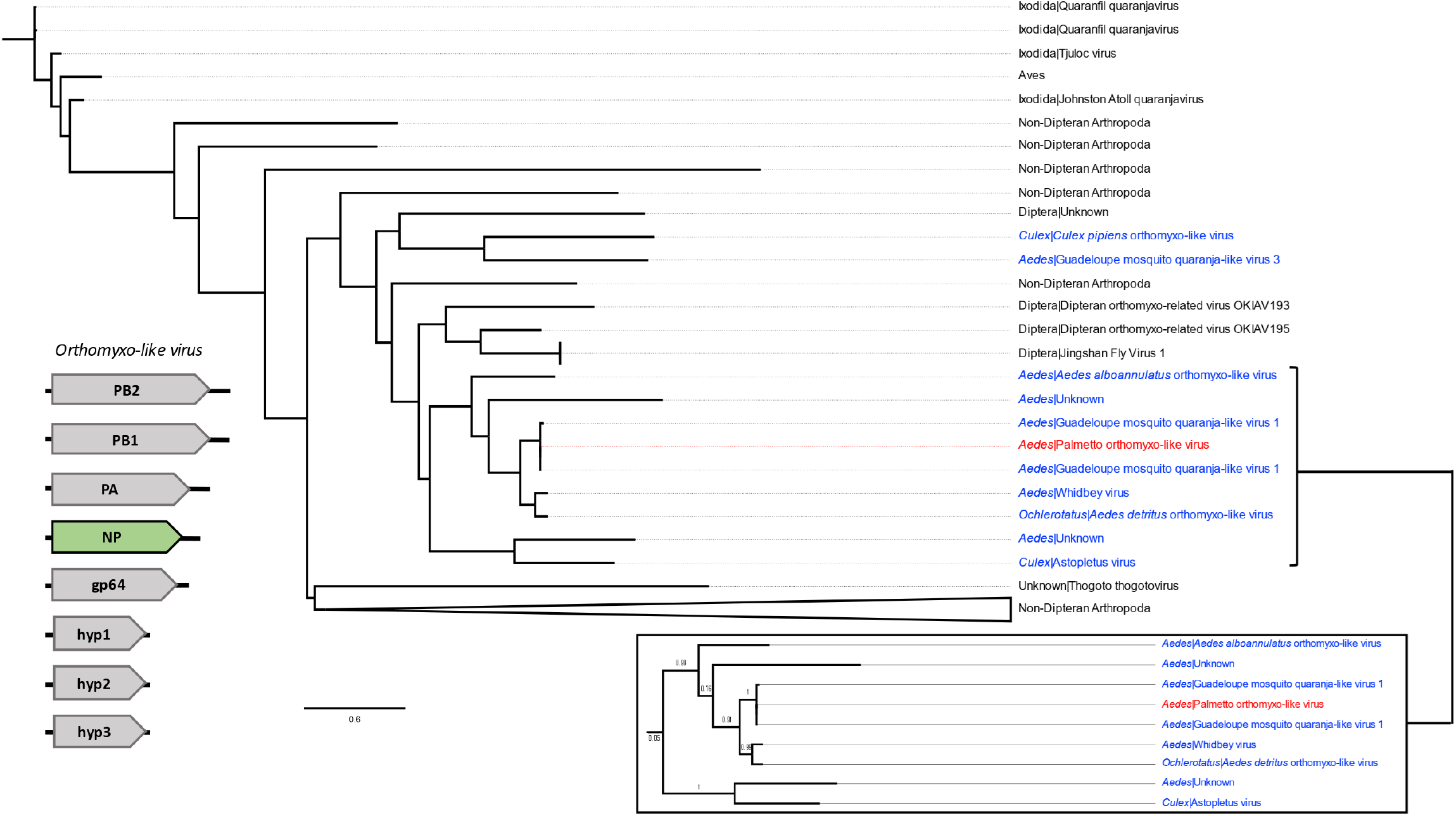
Maximum likelihood tree of orthomyxo-like nucleoprotein (NP) virus sequences (using an LG+G+F model) based on amino acid sequences. The tree is representative of 63 sequences spanning 932 sites and 123 branches. Mosquito-derived virus sequences are colored in blue, while the Palmetto sequence is in red. Virus names are appended to all sequences derived from Diptera or Ixodida. The generalized viral genomic structure is to the right of the tree, with the segment analyzed in color (green polygon represents capsid/nucleoprotein sequence). The numbers on each branch within the inset denote branch support lengths. Scale bar is representative of 0.6 amino acid substitutions per site.

### Reverse transcription PCR confirmation supports virus vertical transmission

To assess vertical transmission success of these three ISVs, reverse transcription PCR with primers designed from metagenomic data for the Palmetto toti-, partiti- and orthomyxo-like viruses (Table S1) was used to re-screen RNA remaining from the original G_0_ pools, as well as RNA from ovaries of G_7_ females and from G_7_ males. This approach yielded amplicons for the toti-like virus capsid gene, the partiti-like virus RdRp gene, and the orthomyxo-like virus PA gene (Figure 9). Amplicons were sequenced and the presence of all three viruses was confirmed in both G_0_ and G_7_ ovary pools (Figure 9 A, C, E). The Palmetto orthomyxo-like virus was present in all G_7_ adult-male pools (Figure 9 B, D, F), while the Palmetto partiti-like virus was found only in a subset of these samples, occurring in 3 of 5 pools. Intriguingly, no males were positive for Palmetto toti-like virus. Although the occurrence and frequency of environmental acquisition and venereal transmission of these viruses remain open questions, the ovary-positive data and presence across multiple generations suggests viral persistence in the population by vertical transmission.

**Figure 9.**
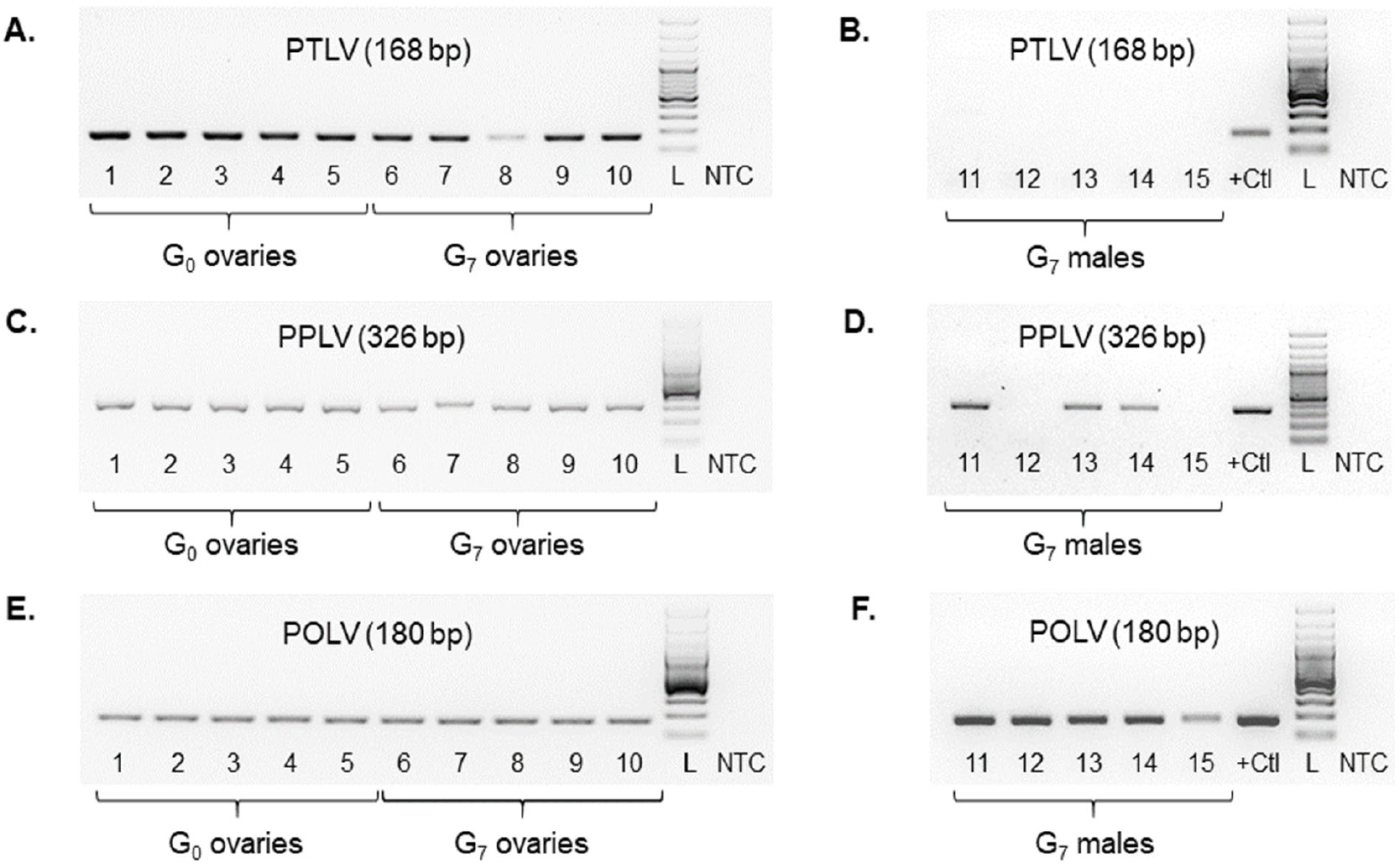
Agarose gel images showing results of RT-PCR screens for Palmetto toti-like virus (PTLV) in ovaries (A) and males (B), Palmetto partiti-like virus (PPLV) in ovaries (C) and males (D), and Palmetto orthomyxo-like virus (POLV) in ovaries (E) and males (F). For each primer set (Table S1), 1 – 5 denote P1 – P5 ovary pools from G_0_ females, 6 – 10 denote ovary pools from G_7_ females, and 11 – 15 denote pools of males. For each set of reactions, NTC is the no template control, L denotes 100-bp ladder, and +Ctl is the positive control for male assays, which consisted of a previously positive ovary pool.

### Viral gene phylogenetics

Phylogenetics on virus gene segments was completed to investigate the evolutionary relationships of each of ISVs to infer their ecological history. Database mining and protein alignments resulted in 25 sequences spanning 1008 sites for the toti-like capsid tree, 101 sequences (1032 sites) for the toti-like RdRp tree, 101 sequences (447 sites) for the partiti-like RdRp tree, 78 sequences (1668 sites) for the orthomyxo-like PB2 tree, 101 sequences (1041 sites) for the orthomyxo-like PB1 tree, 101 sequences (1388 sites) for the orthomyxo-like PA tree, and 63 sequences (932 sites) for the orthomyxo-like NP tree. After Smart Model Selection (SMS), five trees resulted in an LG optimal model with Gamma rates (G), invariable sites (I), and empirical frequencies (F) (LG+G+I+F). Log-likelihood ratios for the toti-like trees were - 53142.30 and -23019.63 (RdRp and capsid, respectively), -44320.2 for the partiti-like RdRp tree, and -77638.40 and -94785.91 for the orthomyxo-like PB1 and PA trees, respectively. The orthomyxo-like PB2 and NP trees resulted in a LG+G+F optimal model with log-likelihood ratios of -86514.82 and -49892.72, respectively.

Both the toti-like capsid and RdRp gene trees showed distinct mosquito clusters (Figures 2 and 3) with strong branch support. The Palmetto toti-like virus identified herein clustered with other *Aedes aegypti* toti-like viruses. *Culex* derived toti-like viruses made up a distinct clade in the RdRp tree, while in the capsid tree two of the *Culex*-derived viruses appeared more closely related to viruses from *Ochlerotatus*, a subgenus of *Aedes* according to traditional Culicidae systematics (Wilkerson et al. 2015). Because of better representation of RdRp sequences in databases, we were able to analyze a broader range of totivirus sequences in the RdRp virus tree revealing distant relatedness to totiviruses from numerous non-dipteran arthropods, fungi and oomycota.

In contrast, the partiti-like RdRp phylogenetic tree showed no obvious mosquito clustering, as mosquito-derived partiti-like virus sequences were dispersed throughout the tree (Figure 4). Our Palmetto partiti-like virus formed a small but well-supported clade with other mosquito partiti-like viruses (Atrato partiti-like virus 3 and Broome partiti-like virus), as well as partiti-like viruses from other non-dipteran arthropods, plants, and molluscs. Virus sequences from non-dipteran arthropods, dipterans, and insects made up most of the remaining branches in the tree.

All the orthomyxo-like trees (PB2, PB1, PA, and NP) displayed similar trends to one another, with two distinct mosquito clusters, loosely organized into *Culex-* and *Aedes-*derived viruses, separated by ISVs from primarily hematophagous Dipterans (Figures 5-8). In all the orthomyxo-like gene segment trees, our Palmetto orthomyxo-like virus was part of the well-supported *Aedes*-based mosquito clade, directly clustering with Guadeloupe mosquito quaranja-like virus 1, Whidbey virus, and *Aedes detritus* orthomyxo-like virus.

## Discussion

### Conservation of a vertically transmitted Floridian *Aedes aegypti* ‘core virome’

Many of the viruses detected through our sequence analysis were present in both the field- and lab-derived samples. Of the sequences common to all samples, the most abundant corresponded to *Aedes aegypti* toti-like virus, although a dsRNA virus environmental sample and an unclassified Riboviria virus were also prevalent (Figure 1). The Palmetto toti-like virus identified here across all *A. aegypti* samples showed striking similarity to *Aedes aegypti* toti-like viruses from Guadaloupe and an unnamed virus identified in *A. aegypti* from Thailand, which was first described as dsRNA environmental virus sample. Totiviridae and toti-like virus samples have been found in several mosquito genera with widespread global prevalence, including *Anopheles* in Liberia (Fauver et al. 2016), *Armigeres* in China (Zhai et al. 2010), *Culex* in Belgium (Wang et al. 2020), *Culex* in California (Batson et al. 2021), *Culex* in Australia (Williams et al. 2020), *Culex* in Japan (Isawa et al. 2011), and *Mansonia* in Brazil (de Lara Pinto et al. 2017). Metagenomic analyses of G_0_ Manatee County mosquitoes in 2016 and 2017 also identified a toti-like virus (*Anopheles* totivirus), as well as a dsRNA virus from an environmental sample (Boyles et al. 2020). This is further support that *A. aegypti* mosquitoes with similar genetic backgrounds (here, representative of Florida-based populations) share the same virus families (i.e., ‘core virome’ components), as previously reported (Öhlund et al. 2019, Shi et al. 2019, Wang et al. 2020, Konstantinidis et al. 2021).

### The field *A. aegypti* insect-specific viral landscape is more diverse

Despite the pooled nature of our samples, there were clear differences in the general diversity of the virus communities, as 12 eukaryotic virus taxa were found in the field-derived samples compared to only five in the ORL sample. This greater virome diversity of field-derived samples was also noted in the 2016-2017 Manatee County G_0_ samples (Boyles et al. 2020), as were numerous other partiti-like viruses (Hubei partiti-like viruses 29, 32, 33, 34, Wenling partiti-like virus 2) and two viruses in Orthomyxoviridae (Whidbey virus, *Aedes alboanulatus* orthomyxo-like virus). The Palmetto partiti-like virus (Partitiviridae) and the four taxa belonging to the family Orthomyxoviridae (Guadeloupe mosquito quaranja-like virus 1, Whidbey virus, *Aedes alboannulatus* orthomyxo-like virus, and an unclassified Orthomyxoviridae virus) (Figure 1) were solely found in the field samples. These differences likely stem from variation between the field population and lab colony. While the field populations were exposed to diverse environments, various feeding sources, and dynamic larval habitats, the lab ORL *A. aegypti* have been raised for decades under stable, standardized conditions. As such, the contrast in ecological conditions experienced by mosquitoes in the lab versus the field may reflect the viral diversity of their respective environments. Although non-vertical routes of ISV acquisition in mosquitoes (i.e., horizontal transmission) are not well studied, to date the data suggest that these viruses may establish new infections through plant or detritus feeding by larvae, directly from water in the larval environment, from the meconium upon adult eclosion, from fungal or parasitic infection, via parasitoids, through nectar feeding by adults, and from one mosquito to another during mating (Vasilakis and Tesh 2015, Dolja and Koonin 2018, Agboli et al. 2019, Xu et al. 2020). As the ORL strain *A. aegypti* have been raised in laboratory settings for over 60 years (Kuno 2010), these diverse viral acquisition routes would be limited.

### Totiviridae and Partiviridae phylogenies point to shared plant- and fungal-based lineages

The clear mosquito-associated clade present in the toti-virus like sequences (Figures 2 and 3) could represent widespread concurrent viral movement or long host co-evolution (Wang et al. 2020), as is thought to be the case for Bunyaviridae, Flaviviridae, and Rhabdoviridae ISVs, which are assumed to have evolved and diversified alongside their host (Vasilakis and Tesh 2015). Historically, totiviruses were primarily fungal associated, and likely represent a viral taxon with an ancient origin (Dolja and Koonin 2018). This historic basis is mirrored in our toti-like virus RdRp tree (Figure 3), as mosquito-associated sequences matched to dipteran and non-dipteran arthropod virus RdRps, followed by distant clades of fungal toti-like virus RdRps. The jump of these toti-like viruses into invertebrate systems is likely due to horizontal virus transfer (Dolja and Koonin 2018), and may have occurred within the mosquito itself, as fungi are common inhabitants of the mosquito microbiome. Similarly, the Partitiviridae and partiti-like viruses were historically associated with plants and fungi but have increased in prevalence among arthropods (Faizah et al. 2020). Partitiviridae and partiti-like viruses have been found across numerous mosquito genera (*Culex, Culiseta, Coquilettidia, Anopheles, Aedes*) with a nearly global distribution (North and South America, Africa, Asia, and Europe) (Öhlund et al. 2019, Shi et al. 2019, Wang et al. 2020, Konstantinidis et al. 2021). The low congruence between viral phylogeny and host-range for these partiti-like viruses may suggest a recent host-switching event (Grubaugh et al. 2016, Dolja and Koonin 2018), as the Palmetto partiti-like virus was highly divergent (<80% similarity) from other known partiti-like viruses and likely represents a novel virus. Cross-species transmission (i.e., horizontal virus transfer) is clearly common throughout the ISV landscape (Shi et al. 2018) and has likely been a major factor in the evolutionary history of the partiti-like virus described herein (Figure 4).

### Orthomyxoviridae phylogenies show a distinct hematophagous arthropod clade with known human pathogenic members

The phylogenies for the Orthomyxoviridae genome segments displayed similar trends, with viruses from non-hematophagous, non-dipteran arthropods as ancestral to those in mosquito-enriched clades in all four trees (Figures 5-8). Known vertebrate pathogenic viruses formed distinct and separate sub-clades from the mosquito and hematophagous dipteran viruses. The former largely included tick-vectored vertebrate viruses such as Bourbon virus, Johnston Atoll quaranjavirus, and Batken virus, among others. The latter was sub-differentiated loosely into *Aedes* and *Culex* ISV sub-clades, with our Palmetto orthomyxo-like virus falling into the *Aedes* clade. Our Palmetto orthomyxo-like virus appears to be nearly identical to the Guadeloupe mosquito quaranja-like virus 1 (GMQLV1) first identified in *A. aegypti* from the Caribbean by Shi et al. (2019), and as such, is likely a member of the genus *Quaranjavirus*. This genus was only recently described (Presti et al. 2009), and genomes of its viruses likely consist of eight negative-sense, single-stranded RNA segments (Batson et al. 2021) (Figures 5-8). Genome fragments recovered from our data matched to genes on four segments. Most known *Quaranjavirus* members differ from their influenza relatives via their surface glycoprotein (gp64), which has similarity to *Baculoviridae* members, and is hypothesized to have been the catalyst for virus entry and fusion in ticks (Allison et al. 2015). These quaranjaviruses have demonstrated horizontal transmission cycles akin to arboviruses between ticks and tropical and subtropical birds (Presti et al. 2009), which is likely why viruses from birds and ticks appear as distant relatives to the Palmetto orthomyxo-like virus in our gene trees (Figures 5-8). Interestingly, these tick-vectored quaranjaviruses showed a lack of replication in *Aedes albopictus* C6/36 cells (Presti et al. 2009, Allison et al. 2015), suggesting that host specificity may be more strictly defined for quaranjaviruses (i.e., less amenable to host switching). In turn, this supports the notion that ISVs vary in their efficiency of vertical transmission, and hence, their capacity to be maintained in a host population by this mechanism over time. The phylogenies based on gene segments for the orthomyxo-like viruses suggest high transmission fidelity, while the partiti-like virus tree suggests greater amenability to host switching. Such patterns may provide clues to ISV ecology, including their degree of specialization and relative ability to use environmental routes to infect mosquitoes horizontally.

### Palmetto toti-like, partiti-like and orthomyxo-like insect specific viruses are vertically maintained in *A. aegypti*

Vertical maintenance of all three Palmetto insect specific viruses identified herein is likely occurring, as we were able to detect each virus across *A. aegypti* generations post-field collection through multiple generations (G_0_ and G_7_) via RT-PCR (Figure 9), a trend noted with numerous ISVs (Lutomiah et al. 2007, Bolling et al. 2011, Haddow et al. 2013, Contreras-Gutierrez et al. 2017, Frangeul et al. 2020). As all ovary pools were positive in both G_0_ and G_7_ generations for all three viruses (Figure 9A, C, E), transovarial transmission was likely the primary route of vertical transmission. As males are known to carry a wealth of ISVs (Frangeul et al. 2020), we also tested pools of adult males for each of our three viruses. Male pools were consistently positive for Palmetto orthomyxo-like virus, positive in the majority of pools for Palmetto partiti-like virus, and notably negative for Palmetto toti-like virus (Figure 9B, D, F). These discrepancies in positivity may suggest differences in transstadial transmission between the sexes during development, which could arise from sex-specific tissue tropisms as adult structures form in the pupal stage. Moreover, testes might display specificity for certain viruses in their small RNA machinery, leading to a more efficient antiviral response (Frangeul et al. 2020). Alternatively, since the toti-like virus was also discovered in our lab-adapted colony (ORL), it may have been acquired during blood feeding. Both G_0_ and G_7_ females were fed on chickens to enable egg production and oviposition prior to ovary dissection. This is the only environmental factor in our study that differs between the sexes, but because totiviruses are not common in domestic birds, we find this route unlikely. For the two viruses present in males, venereal transmission may complement vertical transmission as a secondary maintenance mechanism for ISV persistence, as has been described for bunyaviruses and alphaviruses in other mosquito species (Ovenden and Mahon 1984, Schopen et al. 1991).

### Metagenomic sequencing utility and relevance

Many of the virus assignments for our *A. aegypti* (Palmetto) mosquitoes matched to unclassified Riboviria lacking further taxonomic resolution, a trend noted in other publications (Öhlund et al. 2019, Shi et al. 2019, Williams et al. 2020). This is likely due to limitations imposed by incomplete virus databases and augmented by the difficulty of recovering segmented virus genomes from pooled samples with high virus similarity (Shi et al. 2019, Batson et al. 2021). As more studies examine viral communities and increase the overall database size, taxonomic classification of this metagenomic ‘viral dark matter’ should drastically improve (Batson et al. 2021).

Despite the presence of DENV-4 in the abdomens of Manatee County mosquitoes in 2016 and 2017 (Boyles et al. 2020), we did not identify any circulating human pathogenic viruses in the ovaries of our field-derived 2018 Manatee County mosquitoes from Palmetto. The effects of the virome, particularly from the presence of various combinations of viruses, are largely still unknown. Studies have shown variation in vector competence with ISV infection (Bolling et al. 2012, Zhang et al. 2017, Baidaliuk et al. 2019) and a positive association between ISV infection and human pathogen infection in mosquitoes (Newman et al. 2011). However, these results do not hold true in all systems (Kent et al. 2010, Crockett et al. 2012). Furthermore, although potential ISV-arbovirus interaction modes, such as competitive inhibition and superinfection exclusion, have been proposed (Vasilakis and Tesh 2015, Roundy et al. 2016,), mechanistic data regarding these interactions, especially on a community scale (i.e., the whole microbiome within a mosquito), are severely lacking. ISV infection in mosquitoes is widely assumed to be commensal (Hall et al. 2016), although examples of ISV-insect interactions with outcomes that depend on the biological context can be found in the literature. For example, a partiti-like virus infection in the fall armyworm (*Spodoptera frugiperda*) causes negative fitness effects in *S. frugiperda* but renders the caterpillar more resistant to a pathogenic nucleopolyhedrovirus (Xu et al. 2020). Furthermore, the low abundance of some of these presumed ISVs and high co-appearance with fungal pathogens may mean that some ISVs are more likely associated with infecting fungi than the mosquito itself (Shi et al. 2017). It remains unclear if similar symbioses are occurring or could occur with the three ISVs studied here.

## Conclusions

Our data suggest that there is a core set of vertically transmitted ISVs in Palmetto *Aedes aegypti* representing at least three virus families: Partitiviridae, Totiviridae and Orthomyxoviridae. The toti-like virus was also prevalent in the ovaries of *A. aegypti* derived from a colony long maintained in the laboratory. We were able to confirm that vertical maintenance of three of these viruses is likely occurring via transovarial transmission, as the Palmetto partiti-, toti-and orthomyxo-like viruses were present in all ovary pools spanning seven generations. Although the orthomyxo-like virus was also present in all male pools, occurrence of the partiti-like virus was less consistent among males, and the toti-like virus was absent from male pools altogether, suggesting potential variation in sex-specificity among ISVs. The striking differences in distant host taxa between the toti-like and partiti-like virus trees compared to the orthomyxo-like virus tree may suggest disparate routes of viral evolution or acquisition (e.g., plant/fungi-based versus vertebrate-based). However, because these data were obtained from pools of ovaries and males, we lack the individual-level resolution to determine ISV prevalence in the population or colony, assess the variability of ISV community composition among individuals, or assess within-mosquito virus abundance. Furthermore, despite previous detection and full genome sequencing of DENV-4 in G_0_ Manatee County *A. aegypti* collected in 2016 and 2017 (Boyles et al. 2020), we did not detect human pathogenic viruses in our metagenomic screen. Nevertheless, this study provides ecological baseline data for ISV occurrence and vertical transmission for a natural, endemic vector species in a state with rising local DENV transmission. Further research on ISV presence/persistence in mosquitoes and their impact on the suite of biological parameters that determine vectorial capacity could help contextualize the role ISVs play in arboviral transmission.

## Materials and Methods

### Mosquito Collections and Sample Preparation

Mosquito eggs were sampled from Palmetto, Florida, in July and August of 2018 in collaboration with the Manatee County Mosquito Control District. Fifteen ovijars were placed throughout a peri-urban area of approximately 1 square kilometer, and ovijar locations were the same as those previously described to sample *Aedes* spp. eggs in 2016-2017 (Boyles et al. 2020) (Figure 10). Egg papers were returned to the insectary at the Florida Medical Entomology Laboratory (FMEL), hatched in distilled water, reared to adulthood, and sorted by species into separate cages by ovijar. Eggs from four ovijars yielded sufficient *A. aegypti* adults for successful mating. On days 5-7 post-emergence, females (G_0_) were fed on chickens (IACUC protocol 201807682) and three days later transferred individually to 50 mL ovicages lined with moist seed germination paper. Gravid mosquitoes were allowed to oviposit over a two-day period, and eggs were dried and stored separately. Upon egg collection, each female was cold anesthetized on ice, surface sterilized in 70% ethanol, rinsed twice with sterile PBS, and then dissected one at a time to remove pairs of ovaries. Following dissection, ovaries were immediately transferred to RNAlater solution (Invitrogen, Waltham, MA) and then combined into five pools from 24-42 females depending on ovijar location and date of egg collection. These pools were then stored at -80°C until processed for RNA. Eggs collected from these mosquitoes were used to establish a field-derived colony from Palmetto. After seven generations of maintenance in the laboratory, ovaries were dissected from 50 G_7_ *A. aegypti* females as described above, combined into five pools of ten pairs, and stored at -80°C in RNAlater. Similarly, 15 G_7_ *A. aegypti* males were collected from the same cage as the females, killed by freezing at -20°C for 15 minutes, and stored as five pools of three individuals in RNAlater at -80°C. These same methods were used on the ORL *A. aegypti*.

**Figure 10.**
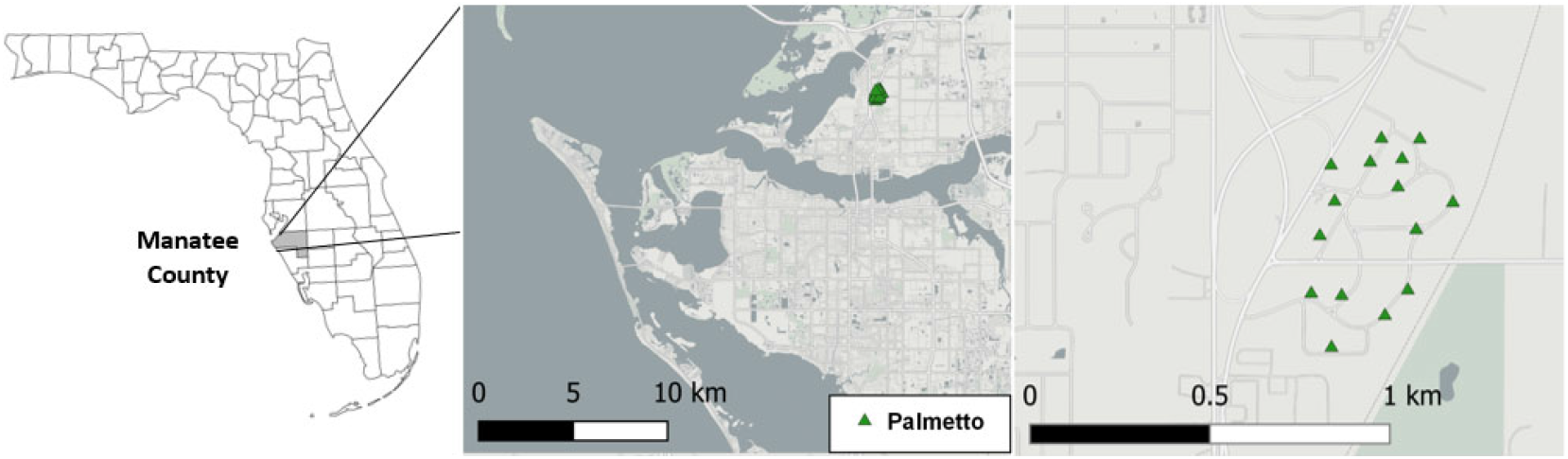
Locations of ovitraps (green triangles) throughout Palmetto, Manatee County, FL. Ovitraps covered approximately 1 square kilometer in a peri-urban (i.e., residential) environment.

Prior to RNA extraction, ovary pools from G_0_ females were thawed on ice, and RNAlater was removed. Pools were homogenized manually in 0.2 mL of TRIzol reagent (Invitrogen, Waltham, MA) using a sterilized pestle. The volume of TRIzol was then brought up to 1.0 mL for each pool, and RNA was extracted following the manufacturer’s instructions, followed by treatment with TURBO DNase (Invitrogen, Waltham, MA). RNA samples were shipped on dry ice to Novogene, where RNA metagenomic libraries were prepared using the NEBNext Ultra II Directional RNA Library Prep Kit for Illumina (New England Biolabs, Ipswich, MA). Libraries were pair-end sequenced (2×150 bp) using the Illumina HiSeq 4000 platform (Figure 11A). RNA from G_7_ males and from ovaries dissected from G_7_ females was extracted as described above and used to investigate vertical transmission by reverse transcription PCR as described below (Figure 11B)

**Figure 11.**
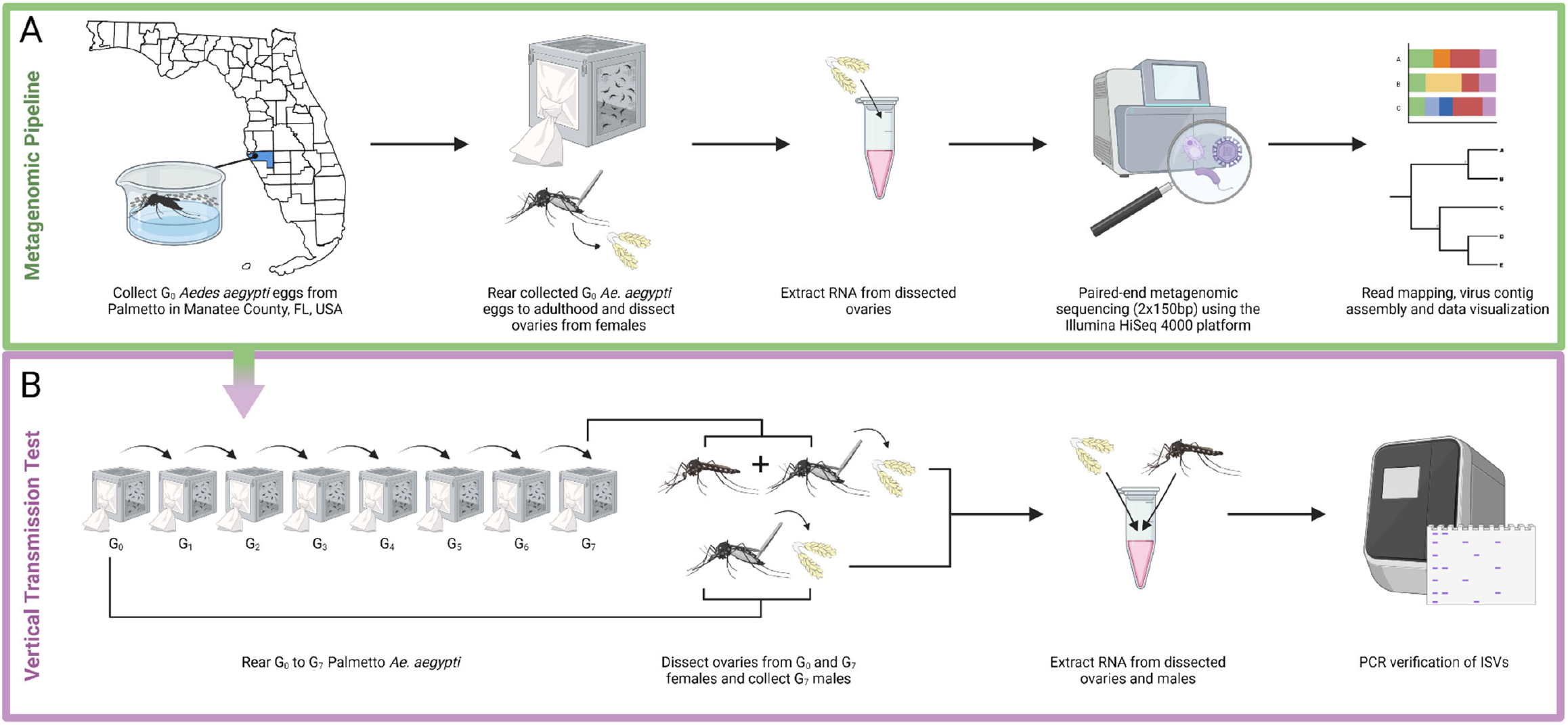
Workflow diagram illustrating the overarching study design for the (A) metagenomic sequencing and (B) vertical transmission assays. In brief, an RNA metagenomic pipeline was employed to discover the insect specific viruses present in the ovaries of *Aedes aegypti* from Palmetto, in Manatee County, FL. Identified ISVs were confirmed via RT-PCR in the G_0_ and G_7_ generations. Figure created with BioRender.com.

### Bioinformatic Analyses of Read level Metagenomic Data

Sequence contaminants and adapters were removed using BBduk v. 38.86 (https://sourceforge.net/projects/bbmap/). *Aedes aegypti* reads were mapped to the *A. aegypti* genome (version AaegL5) and removed using BBmap. Local similarity searches were performed against the National Center for Biotechnology Information’s non-redundant protein database (NCBI nr) (downloaded 01/21) (NCBI Resource Coordinators 2018) using DIAMOND v 2.0.7 (Buchfink et al. 2015). MEGAN6 v 6.19.2 (Huson et al. 2007), was used to assign reads to each lowest common ancestor (LCA) using the default naive LCA settings (in read count mode), mapping against NCBI nr (megan-map-Jan2021.db.zip from the MEGAN6 website (https://software-ab.informatik.uni-tuebingen.de/download/megan6)).

### Viral Genome Assembly

All *A. aegypti* unmapped read files were merged, and contigs were assembled *de novo* using SPAdes v 3.14.1 (Bankevich et al. 2012) in metagenomics mode using merged and unmerged reads. Reads found to align to Atrato partiti-like virus 3 were separately parsed into contigs using the same pipeline (sub-divided due to low similarity with known database matches). Local similarity searches were performed against NCBI nr (NCBI Resource Coordinators, 2018) using DIAMOND v 2.0.7 (Buchfink et al. 2015). MEGAN6 v 6.19.2 (Huson et al. 2007), was used to assign each LCA (using the long-read import option), and to visualize and extract viral contigs. NCBI’s open reading frame (ORF) finder (https://www.ncbi.nlm.nih.gov/orffinder/) was used to identify ORFs in each assembled viral contig. ORFs >300 nucleotides in length were searched using BLAST (Zhang et al. 2000) against the nr protein sequence database through NCBI.

### Reverse Transcription PCR

For a subset of ISVs identified through metagenomics, mapped viral reads from each sample were aligned to create consensus sequences using the Velvet Optimiser (v2.2.6) (Zerbino et al. 2008) to enable primer design for reverse transcription PCR to confirm viral RNA in the original pools and screen tissues from subsequent generations. Consensus sequences were then searched against the NCBI nr database (NCBI Resource Coordinators 2018) using Blastx to confirm the virus and protein match. Primers were designed using Primer3 with default parameters to yield amplicons ranging from 150 – 500 bp (Table S1).

For reverse transcription PCR, we performed a two-step procedure: cDNA first-strand synthesis using an RNA template followed by traditional PCR. For cDNA synthesis, 5 µg of total RNA from each ovary pool (field- and colony-derived) was used to generate cDNA with the RevertAid First Strand cDNA Synthesis kit (Thermo Scientific, Waltham, MA) following the manufacturer’s instructions for random hexamers. PCR amplifications were performed on each pool with all primer sets in 25 µl reactions, each containing dNTPs at a concentration of 0.2 mM, primers at a concentration of 0.2 µM, 1.0 units of Taq DNA polymerase (DreamTaq, Thermo Scientific, Waltham, MA), 1x Taq polymerase buffer, and 1 µl of cDNA from the first-strand synthesis reaction diluted 1:1 with ultrapure water. All PCR amplifications were performed with an initial melt step of 95°C for 5 minutes followed by 35 cycles of 95°C for 30 seconds, 60°C for 30 seconds, and 72°C for 30 seconds, followed by a final extension step of 72°C for 10 minutes.

Amplicons were electrophoresed on 2% agarose gels, stained with GelRed nucleic acid stain (Biotium, Fremont, CA), and visualized on a gel documentation system (Azure Biosystems c200). For samples yielding a PCR amplicon, products were purified using the GeneJET PCR Purification Kit (Thermo Scientific, Waltham, MA). Purified amplicons were ligated into plasmids using the CloneJET PCR Cloning Kit (Thermo Scientific, Waltham, MA) and then incubated with chemically competent *E. coli* (DH5α) (New England Biolabs, Ipswich, MA) following the manufacturer’s instructions. Transformed *E. coli* were grown overnight on selective LB agar plates with carbenicillin (100 µg/mL) at 37°C. Colonies carrying the plasmid and amplicon were transferred to liquid LB media with carbenicillin (100 µg/mL) and grown overnight shaking at 220 RPM at 37°C. Cultures were harvested by centrifugation in microcentrifuge tubes at 17,000 x g for 5 minutes, and plasmid DNA was extracted using the GeneJET plasmid miniprep kit (Thermo Scientific, Waltham, MA). Plasmid DNA was sent to Eurofins Genomics for sequencing using a plasmid-specific primer included with the CloneJET kit.

### Viral Gene Phylogenetics

Alignments were created from resultant viral RdRp and capsid sequence BLAST results using MUSCLE (Edgar, 2004) in MEGA X 10.0.3 (Kumar et al. 2018) (Supplementary Files 1-7). PhylML v 3.0 (Guindon et al. 2010) and Smart Model Selection (SMS) (Lefort et al. 2017) were used to create an optimized maximum likelihood (ML) phylogenetic tree for each alignment. Branch supports were computed by an approximate likelihood ratio test (aLRT) with SH-like support as implemented in PhyML. Trees were annotated and colored in FigTree v1.4.4 (https://github.com/rambaut/figtree/releases).

## Data Availability

Cleaned and trimmed sequence files have been deposited in NCBI as a BioProject and will be made publicly available upon publication.

## Acknowledgments

We would like to thank X. Wang and T. Stenn for assistance with mosquito feeding and colony maintenance and J. Crosby for animal husbandry. A fellowship for J. Bozic and an internship for J. Carrillo were supported by funds through the CDC Southeastern Center of Excellence in Vector-Borne Diseases (U01CK000510).

## Competing Interests

The authors declare no competing (financial or non-financial) interests.

## Notes

### Competing Interest Statement

The authors have declared no competing interest.

