## Supplemental Table 1 for "Intrinsic variation in the vertically transmitted insect-specific core virome of *Aedes aegypti*"

**Table S1.** Primers used in reverse-transcription PCR to screen RNA pools derived from *Ae. aegypti* ovaries ($G_0$ and $G_7$) and males.

| Virus | Direction | Sequence (5' - 3') | Amplicon Length |
|---|---|---|---|
| Palmetto toti-like virus | Forward | CCATCTGAAAGAGAGCAGAC | 169 bp |
| | Reverse | TCGGTCGAAATGTGTGGTAA | |
| Palmetto partiti-like virus | Forward | TTGCACTCCCTTTGATCGAC | 326 bp |
| | Reverse | CTTCCACCTTTCGCCGTTAC | |
| Palmetto orthomyxo-like virus | Forward | AACCCATGAGGCGTCAATAC | 180 bp |
| | Reverse | GTTTCCCTTTGGTTCTGCAA | |
